# Steady-state polypeptide transfer from the translocon to the membrane

**DOI:** 10.1101/2023.01.10.523415

**Authors:** Denis G. Knyazev, Mirjam Schaur, Roland Kuttner, Christine Siligan, Nikolaus Gössweiner-Mohr, Nora Hagleitner-Ertugrul, Peter Pohl

## Abstract

In concert with irreversible non-equilibrium peptide translation by the ribosome, the nascent polypeptide chain may integrate into the membrane or translocate to the other side of the membrane, facilitated by the conserved protein translocation channel SecYEG in bacteria and Sec61 in eukaryotes. Assuming equilibrium for the decision processes yielded the biological hydrophobicity scale, reflecting free-energy differences ΔG between the pore interior and membrane. Yet kinetic effects and molecular dynamic simulations suggested that a nascent polypeptide could not sample the two separate environments a sufficient number of times for partitioning in equilibrium. Here we tested the hypothesis employing purified and reconstituted SecYEG harboring a stalled ribosome nascent chain (RNC). The SecYEG-RNC complex was open in a de-energized membrane, allowing ion flow. Application of a membrane potential closed the channel if nascent chain hydrophobicity permitted membrane integration. Taking the ratio of steady-state to initial ion conductances as a measure of nascent chain hydrophobicity, we found ΔG for KvAP’s voltage sensor (4^th^ helix harboring four arginines) and FtsQ’s transmembrane helix to be equal to 0.3 and –2.1 kcal/mol, respectively. Thus, ΔG observed in our minimalistic system agrees very well with the position-dependent amino acid contribution of the biological hydrophobicity scale. Characteristic sampling times of ~2 s appear sufficient to reach a steady state for a ~20 amino acid-long segment invalidating the hypothesis of insufficient sampling.

## Introduction

The processes of (i) protein insertion into the membrane, (ii) membrane-protein folding, (iii) protein-protein interactions within membranes, and (iv) membrane-protein conformational changes are interrelated (1, 2). Despite their fundamental importance, many aspects of the underlying energetic expenditure thus far remained enigmatic (3–5). As a result, many different hydrophobicity scales have been proposed (6–8). The comparison of two specific amino acid hydrophobicity scales provides an example: Measurements of the apparent free energy ΔG of transferring an amino acid from the aqueous solution into a hydrophobic solvent (octanol) gave rise to the physicochemical hydrophobicity scale (9). It indicates insertion costs of ~3.6 kcal/mol for the most hydrophilic amino acid and a gain in free energy of 2 kcal/mol for the most hydrophobic amino acid. The so-called biological hydrophobicity scale spans a smaller range of only 4 kcal/mol. It was derived in dog pancreas rough microsomes by determining the membrane insertion probability of an alanine-leucine helix doped with the amino acid of interest in its middle. The most hydrophobic amino acid yielded a gain in energy of only ~0.6 kcal/mol (10). Most strikingly, the rank order in the two scales is not the same. The difference between the two scales is mainly attributed to their unequal reference points: bulk water versus aqueous translocon interior.

The biological scale reflects the free energy ΔG_b_ for the amino acid residue transfer from the translocon to the membrane (10). Predicting ΔG_b_ is difficult, partly because the physical properties of the intraluminal water molecules and bulk water are different (11). The observation is consistent with the standard rule that water molecules regain their bulk properties starting from a distance of about five water molecules from the channel wall (12). Due to differences in the hydrophobicity of the pore lining residues, variable channel diameter, and membrane inhomogeneities, ΔG_b_ is position-dependent (13). Moreover, ΔG_b_ for the same amino acid also depends on the translocon, e.g., there are quantitative differences between Sec61-mediated insertion of transmembrane segments into the membrane of the endoplasmic reticulum in Saccharomyces cerevisiae and mammalian cells (14). Moreover, mutations in the hydrophobic core of the Sec61 translocon alter ΔG_b_ (15).

The endpoints also differ for the two hydrophobicity scales. The physicochemical scale reflects the free transfer energy, ΔG_pc_, into octanol (oil, hexadecane) rather than into the membrane interior. This simplification is less of a problem than the mismatched starting points because it has long been established that ΔG_pc_ for the transfer from water into the organic solvent scales with the energetic expense for membrane partitioning. The interrelation between ΔG_pc_ and membrane permeability is known as Overton’s rule. This rule has withstood the test of time (16). It allows for predicting membrane permeability, especially when substituting the one (oil) slab membrane model for more sophisticated multi-slab models (17–19).

Calculating ΔG_pc_ from ΔG_b_ and vice versa appears to be very difficult as it would require many corrections. The most important ones concern the specific amino acid location, the hydrophobicity of the organic solvent, and the free energy of the transfer from bulk water to the translocon. The example of the voltage sensor of a potassium channel, more precisely of KvAP’s fourth transmembrane helix (TM4) shows the complexity of the situation. The structure of the KvAP channel promised an understanding of excitability; hence membrane insertion and movement of KvAP’s voltage sensor got special attention (20, 21). The initially proposed movement of its four gating charges through the hydrophobic membrane core (22) seemed to be at odds with the high Born energy required for the purpose (23).

Indeed, if calculated as the sum of the amino acids’ individual (position-wise uncorrected) ΔG_b_ values, one finds the free energy ΔG_s_ required for the membrane integration of the 19 amino acid-long segment. ΔG_s_ for TM4 insertion amounts to ~3.9 kcal/mol (10). Such ΔG_s_ corresponds to an insertion probability P = 0.1%. In contrast, the physicochemical scale predicts P = 99.9 %. However, the same experiments that led to the biological hydrophobicity scale demonstrated a TM4 insertion probability P = 37% (corresponding to ΔG_s_~0.3 kcal/mol) into the membrane of the endoplasmic reticulum (24). These experiments gave rise to the position-dependent ΔG_b_ values (13), yet it is unknown whether the prokaryotic translocon processing K_v_AP agrees in its hydrophobicity assessment with eukaryotic Sec61.

Quantitative differences in membrane integration may also arise from variations in the folding propensities of adjacent sequences and chaperone binding (25). The observation raises the question of whether the presumed equilibrium, a pre-requisition for the ΔG_s_ calculation from ΔG_b_ values, is met during protein integration into the membrane. Moreover, the solvation energies per unit area for partitioning of aromatic side chains are a factor 2.5 smaller than found in the physicochemical scale’s settings (biphasic water-oil), suggesting that translocon-to-membrane partitioning may be biased by other effects (26). Moreover, extensive molecular dynamics simulations raised doubts that the nascent chain has sufficient time to sample the different environments of the aqueous translocon pore and the membrane interior (27). They suggested that kinetic effects may govern the membrane insertion process.

Here we used an *in vitro* system that provides the nascent chain with ample time to probe the hydrophobicities of the membrane interior and translocon pore (28). To allow the stalled translocation intermediate to reach a steady state location, we reconstituted purified SecYEG-RNC (ribosome-nascent chain) complexes into planar lipid bilayers. Movement of the stalled nascent chain out of the aqueous channel manifested as a loss of ion channel activity due to channel closure. Electrophysiology showed renewed ion channel activity when the nascent chain returned to the lumen of the bacterial translocon SecYEG.

We monitored the partitioning of two transmembrane helices (TM) with different hydrophobicities: the strongly hydrophobic TM of FtsQ and the weakly hydrophobic TM4 of KvAP. To quantify the impact of positive charges on the partitioning, we also varied TM4’s number of arginine residues. The thus obtained ΔG_s_ for KvAP’s TM4 is in perfect agreement with the ΔG_s_ from experiments in dog pancreas rough microsomes (24), implying that sampling limitations are unlikely to bias polypeptide membrane insertion.

## Materials and Methods

### Electrophysiological measurements

Ag/AgCl reference electrodes were immersed into the buffer solutions on both sides of the planar bilayers (Fig. 1A). The command electrode of the patch clamp amplifier (model EPC9, HEKA electronics, Germany) localized to the cis compartment, and the ground electrode to the trans compartment. The recording filter for the transmembrane current was a 4-pole Bessel with a −3 dB corner frequency of 0.1 kHz. The raw data were analyzed using the TAC software package (Bruxton Corporation, Seattle, WA). Gaussian filters of 12 Hz were applied to reduce noise. The data were further processed using SigmaPlot (Systat Software Inc., San Jose, CA).

**Figure 1.**
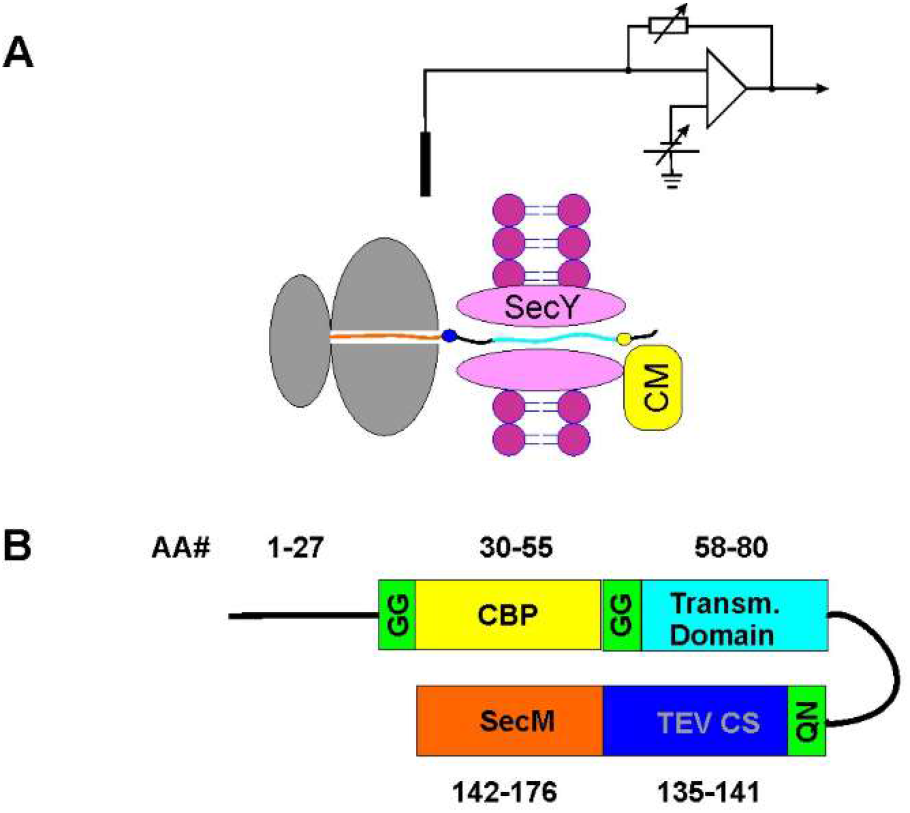
Experimental approach. **A:** Schematic representation of the locked translocation intermediate. Calmodulin (CM, yellow) binds to the calmodulin-binding tag (CBP, yellow) that protrudes into the hypotonic compartment at the *trans* side of the planar lipid bilayer. At the *cis* (hypertonic) side of the bilayer, the ribosome (gray) anchors the transmembrane domain (cyan) of the nascent chain at the SecM stalling sequence (orange). **B:** The nascent chains contain the N-terminal part of FtsQ (black lines), a variable transmembrane segment (Table 1), a SecM stalling sequence (orange), and two affinity tags: TEV cleavage site (blue) for Western blot analysis and CBP. The linking amino acids are depicted in green (see the SI for the exact amino acid sequence).

**Table 1.** Transmembrane domains of RNCs.

| Transmembrane domains of RNCs | TM Sequence |
| --- | --- |
| KvAP 4R (wild type) | GG FRLVR LLRFL RILLI IS DA |
| KvAP 3R | GG FRLVR LLAFL RILLI IS DA |
| KvAP 0R | GG FALVA LLAFL AILLI IS DA |
| FtsQ | GG LFLLT VLTTV LVSGW VV LG |

### SecYEG purification

Purification of SecYEG was performed as previously described (29, 30). Briefly, SecYEG was overexpressed for 3 h in E.coli c43 (DE3) cells from a pBad22 vector and induced with 2 g/l of arabinose. The collected cells were lysed with an Emulsiflex homogenizer (Avestin) in 20 mM Tris (pH 7,5); 300 mM NaCl; 10 % glycerol; supplemented with complete protease inhibitor (Roche). The membrane fraction was pelleted at 100.000 × g and solvated in 1 % (w/v) Dodecyl-malto-pyranoside (DDM, Anatrace). Affinity chromatography with Ni-NTA-Agarose (Peqlab) and size exclusion chromatography were used to improve sample purity.

### Purification of RNC constructs

Affinity purification of the stalled RNC constructs (Fig. 1A) occurred in two steps. First, we used Ni chelating agarose (Quiagen) and, second, calmodulin agarose (Sigma) as previously described (31). Accordingly, the RNCs contained two tags for affinity purification: a 6xHis-tag at the L12 protein of the 50S subunit (32) and a calmodulin-binding peptide (CBP) (Fig.S1-2). All RNCs harbored FtsQ’s 101 N-terminal residues and the SecM stalling motif FXXXXWIXXXXGIRAGP (Fig. 1B). Yet, the constructs differed in their transmembrane domain (see Table 1).

The constructs were expressed for 2 h in JE28 cells containing his-tagged ribosomes by the addition of 2 g/l arabinose. Pelleted cells were lysed in 20 mM Tris (pH 7.6), 10 mM MgCl_2_, 150 mM KCl, and 30 mM NH_4_Cl in a homogenizer at low pressure (15-17 kPsi).

### SecYEG reconstitution into lipid vesicles

SecYEG was reconstituted into E.coli polar lipid extract (Avanti Polar Lipids) vesicles pre-dissolved in deoxy-BigChap (Anatrace) as previously described (33). Biobeads SM2 (Biorad) were added to remove the excess detergent, and the resulting turbid suspension was pelleted at 100.000 × g. The resulting pellet was resuspended and extruded through a 100 nm filter. We used a mass ratio of protein to lipid of 1:100. Reconstitution efficiency was about 3 SecYEG complexes per vesicle, which was established using fluorescence correlation spectroscopy (34). In brief, we labeled purified SecYEG prior to reconstitution with AlexaFluor647 (ThermoFisher Scientific) and compared the number of fluorescent particles in the sample containing proteoliposomes with the number of particles (micelles) after adding 2% octyl glucoside (Sigma) and 3% deoxy big CHAP (Anatrace) to the same sample and correcting for dilution (Fig. S3).

### Locking the translocation intermediates in the reconstituted translocon

Solvent-depleted planar lipid bilayers were formed by folding lipid monolayers from *E.coli* polar lipid extract (Avanti Polar Lipids) in the ~150 μm wide aperture of a PTFE film (Goodfellow).

### SecYEG reconstitution into planar bilayers proceeded via osmotically induced lipid vesicle fusion

About 2 - 5 nM SecYEG-containing vesicles and 10 nM RNCs were added to the hypertonic compartment. It contained 450 mM KCl and 5 mM MgCl_2_ to trigger vesicle fusion with the planar bilayer. The KCl concentration in the hypotonic compartment was limited to 150 mM. Both compartments contained 50 mM K-HEPES and were kept at pH 7.5. Formation of the RNC SecYEG complex resulted in open, i.e., ion-conducting pores. The resulting ion flow maintained hypertonicity in those vesicles that (i) contained the activated translocon and (ii) were docked to the planar bilayer. Water uptake from the hypotonic compartment led to vesicle swelling and fusion. The resulting stepwise rise in membrane conductivity indicated the reconstitution of RNC-SecYEG complexes into the planar bilayer (Fig. 3A). In contrast, vesicles with closed channels, i.e., vesicles in which the RNC-SecYEG complex did not form, are unable to fuse with the planar bilayer (35, 36).

**Figure 2.**
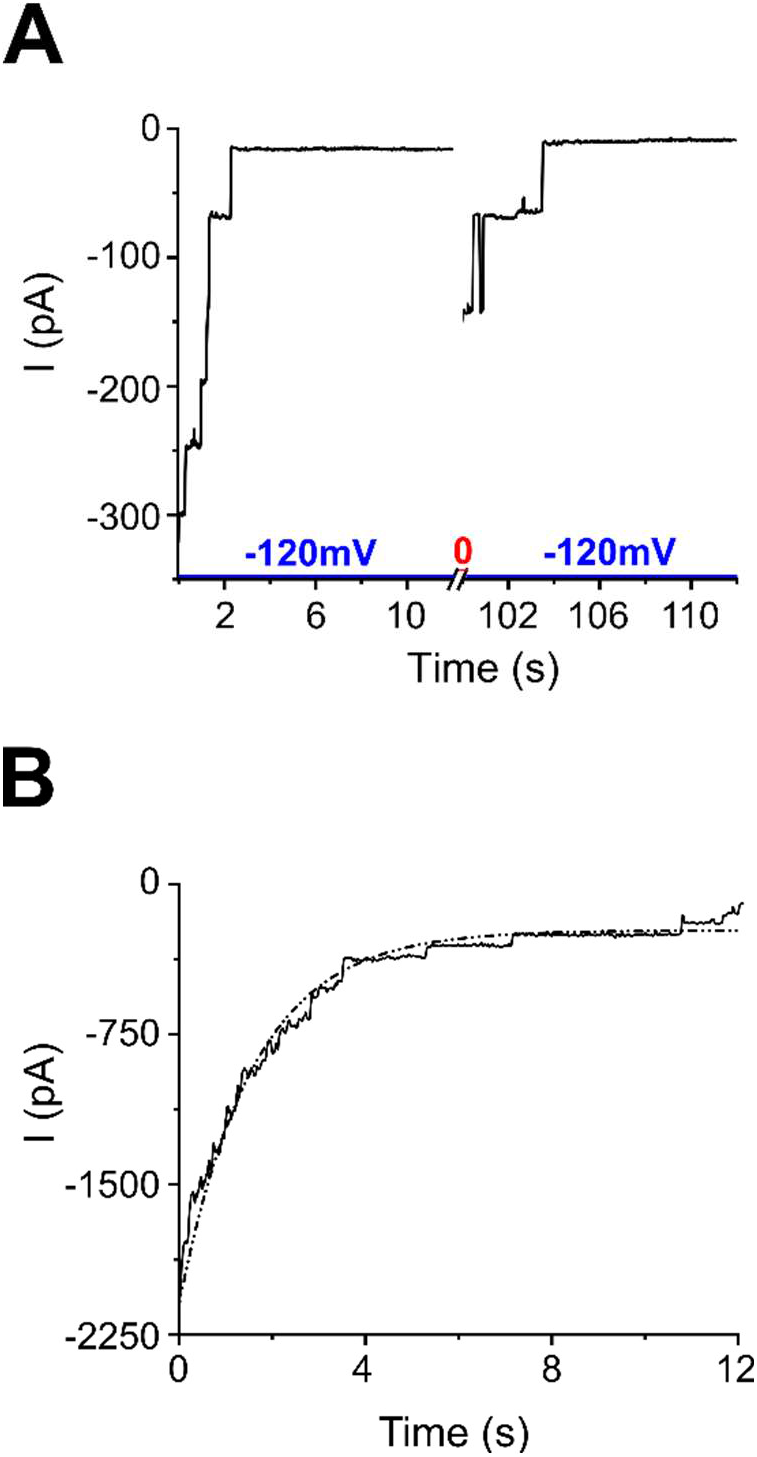
Closing kinetics of translocation intermediates without calmodulin. **A:** Closing of five SecYEG-RNC translocation intermediates with FtsQ-based nascent chain. The voltage protocol consisted of steps −120 mV (blue line), 0 mV (red line), and −120 mV (blue line). Each step lasted 50 s. Five translocation intermediates closed upon application of *ψ* = −120 mV (left part of the graph). After the *ψ* was switched to 0 mV, two of the intermediates reopened, as seen when *ψ* = −120 mV was reapplied (right part of the graph). **B:** Cumulative closing kinetics of SecYEG-RNC complexes with FtsQ-based RNC was built from individual traces comprising 35 translocation intermediates. We fitted the trace with a single-exponential function (dashed dotted line, see Table 2).

**Fig. 3.**
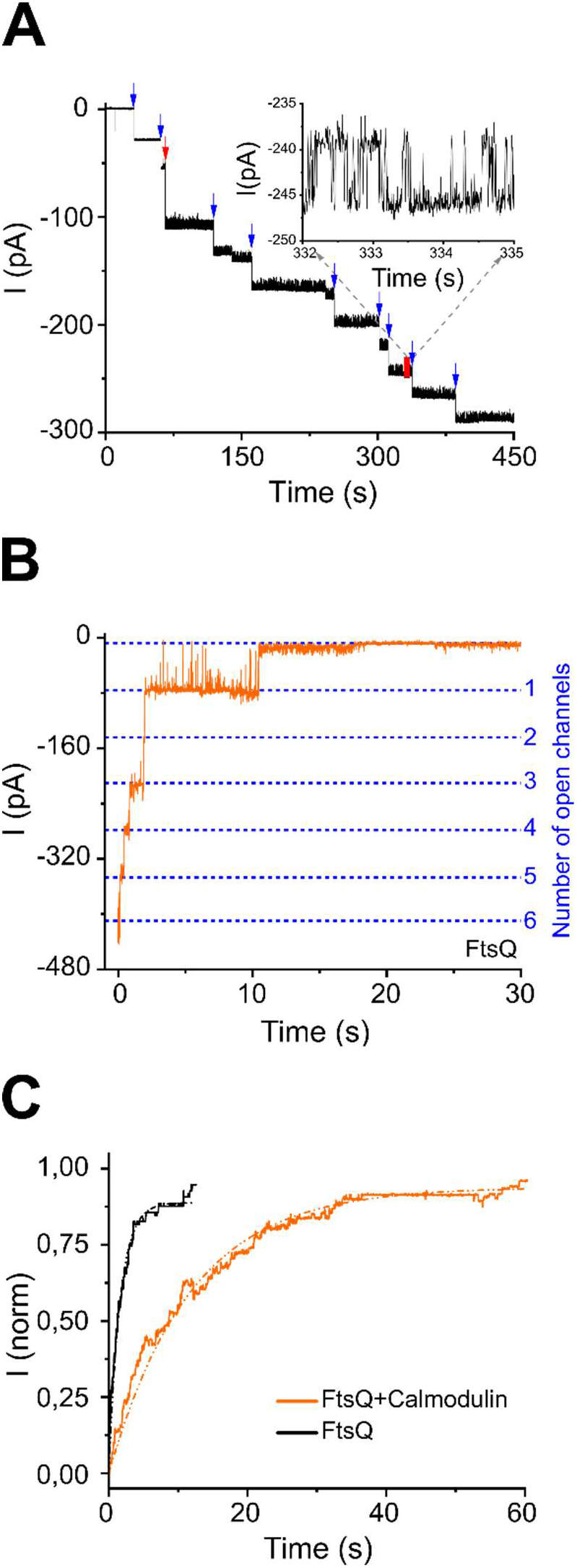
Electrophysiology of SecYEG-RNC(FtsQ). **A.** Electrophysiological record showing the reconstitution of SecYEG-RNCs with a transmembrane segment from FtsQ into the planar bilayer via vesicle fusion. Fusion events of vesicles containing 2 and 4 SecYEG-RNC complexes are indicated by blue and red arrows, respectively. In total, 22 channels were incorporated within 400 s. The inset shows a zoom into the labeled red record region to illustrate the fluctuation of a single pore between the open and closed states. The transmembrane potential *ψ* amounted to −18 mV. All experiments were performed in 50 mM K-HEPES buffer with 5 mM MgCl_2_ (hypertonic compartment) and 1 mM CaCl2 (hypotonic compartment) at pH 7.5. A 450 to 150 mM transmembrane KCl gradient served to facilitate fusion. **B.** SecYEG-RNC(FtsQ) closure events in the presence of calmodulin. The trace shows six translocation intermediates which start closing one by one upon application of a transmembrane potential ψ = −105 mV at t = 0 s. The closings manifest themselves as a stepwise decrease of the ionic current through the membrane and indicate the partitioning of the TMD into the lipid bilayer. **C.** Cumulative trace for SecYEG-RNC(FtsQ) closure. The trace in the absence of calmodulin (see figure 2B) is shown for comparison.

**Table 2.**
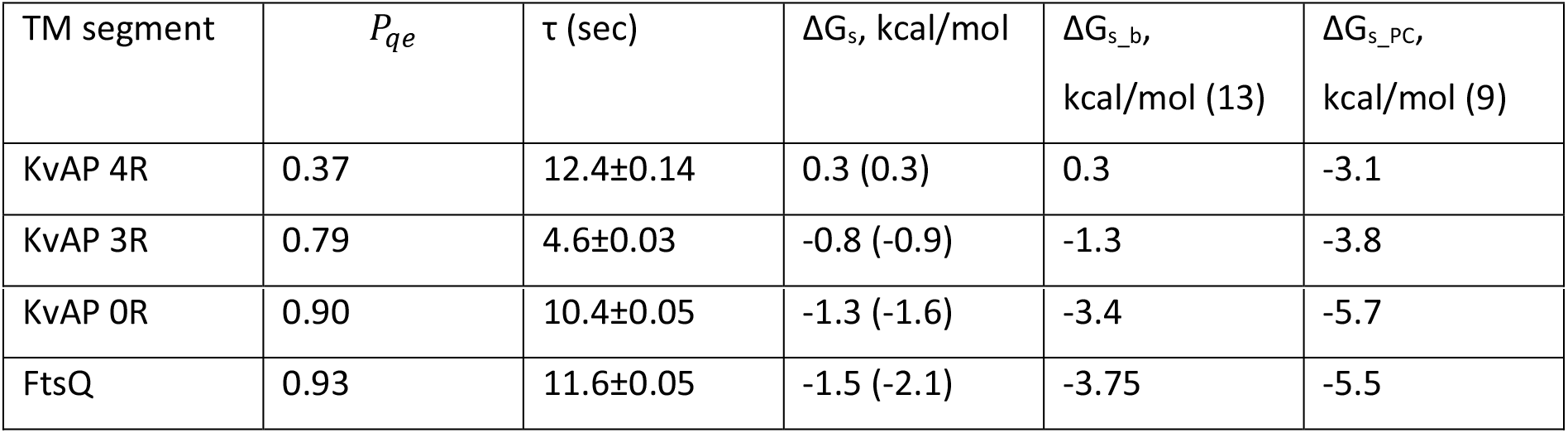
Comparison of experimentally obtained ΔG_s_ with the predictions from physicochemical scale, ΔG_s_PC_, and position-dependent biological scale, ΔG_s_b_.

In most of the experiments, we added 7 μM calmodulin (CM) to the hypotonic compartment to prevent nascent chain backsliding. With the calmodulin bound to the calmodulin-binding sequence at one side and SecM arrested in the ribosome on the other side, the nascent chain was locked. That is, neither retrograde nor forward movements of the nascent chains were possible. They had only one remaining degree of freedom: they could move in and out of the lateral gate. Control measurements confirmed that calmodulin did not cause SecYEG openings in the absence of RNCs.

## Results

### Voltage-driven SecYEG closure triggers nascent chain partitioning into the lipid bilayer from the stalled translocation intermediate

We added SecYEG-containing vesicles and RNCs to the *cis* side (hypertonic side) of the preformed planar bilayer. If added separately, neither SecYEG-containing vesicles nor RNCs induced a discernible current through the planar lipid bilayer. This observation indicates that RNC complexes do not hamper the membrane barrier to ions and that the SecYEG channels are closed in the absence of a ligand. If added together, RNC binding activated vesicular SecYEG channels with an inside-out orientation. Translocons with an inside-in orientation would remain electrically silent since the ribosome binding site was inaccessible. Channel activation enabled the osmotically induced fusion of SecYEG-RNC(FtsQ) containing vesicles with the planar bilayer (Fig. 3A).

The fusion events left the bound ribosomes on the *cis* side of the planar bilayer. Thus, all activated SecYEG channels had the same orientation. They rarely closed at low membrane potentials *ψ*. In agreement with previously reported data, we had to apply near to physiological *ψ* values to close the translocon (28, 30). Applying such *ψ* produced fast stepwise closing events (Fig. 2A, *ψ* = −120 mV, negative polarity at the *cis* side). Closure of the translocon intermediate requires the nascent chain to (i) partition into the lipid bilayer or (ii) slide back into the *cis* compartment and the ribosome to unbind. In the particular recording shown in Fig. 2A, only two out of five channels reopened with the application of −120 mV after a short period of relaxation to 0 mV. To ascertain the dynamics of channel closure, we repeated the experiment shown in figure 2A (left panel) many times. The channel open probability was rather low, so we always observed only a few channels. Adding up all events from different experiments produced Fig. 2B. The y-axis shows the cumulative conductance.

Fitting the conductance with a single exponential function yielded a time constant of 1.6 ± 0.1 s. That is, those of the 19 amino acid-comprising nascent chains that partitioned into the lipid bilayer either already had left the aqueous environment or made that decision at a rate of roughly 10 amino acids per second. Since the translocon closure occurs much faster than empty ribosomes are bound, the latter scenario is the most probable. The rate of 10 amino acids per second is remarkably close to the rate previously observed for nascent chain secretion (37). It is also close to the ribosome’s translation rate of 5 – 10 a.a./s at a high growth rate for yeasts (38) and 15 a.a./s in E. coli (39). Our observation indicates that the rate at which the nascent chain may sample the different environments of the translocation pore and the lipid bilayer suffices to integrate a transmembrane helix in thermodynamic equilibrium. Unfortunately, the possibility of backsliding does not allow us to judge whether the steady state reached in the experimental settings of Fig. 2 mirrors that in a living cell.

### Reconstitution of locked SecYEG-RNC complexes

We performed a new series of experiments where we prevented backsliding by adding CM to the trans-compartment. As in the case without CM, channel activation enabled the osmotically induced fusion of SecYEG-RNC(FtsQ) containing vesicles with the planar bilayer (Fig. 3A). As the FCS experiments indicated the presence of three SecYEG channels per vesicle (Fig. S3), and vesicle fusion inserts all of them at once, we expected to see a correspondingly large increment in planar bilayer conductivity. In line with the prediction, we mostly found fusion events where the incremental current comprised from 2 (blue arrows) to 4 times (red arrow) the current through a unitary SecYEG pore (Fig. 3A).

The nascent chain was locked in a bilayer-spanning position with the ribosome on one side of the membrane and CM bound to the CBP tag on the other side. Consequently, nascent chain movement was restricted to sampling between the aqueous channel and the lipid interior. Also, in these settings, physiological values of the membrane potential closed the channel (−105 mV), forcing the nascent chain to partition into the lipid bilayer (Fig. 3B). Fitting the cumulative conductance curve with the single exponential yielded the characteristic time for channel closure (Fig. 3C). With 11.6 s it was nearly tenfold slower than in the absence of calmodulin. The observed calmodulin effect is in line with the observation that chaperon binding affects membrane helix integration (25). While chaperons with ATPase activity lower the hydrophobicity threshold for membrane integration (25), the binding of a passive ligand merely prolonged the time required to establish a steady state.

We defined the quasi-equilibrium probability *P_qe_* as the ratio between the number *N_p_* of RNCs partitioning into the bilayer and the total number N_total_ of reconstituted SecYEG-RNC complexes:

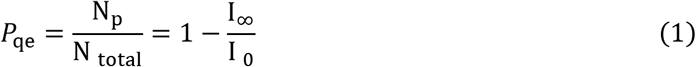

where *I*_0_ and *I*_∞_ are the steady-state ionic currents through the bilayer immediately after applying *ψ* (at time t = 0) and after the steady-state has been reached (at time t = ∞). *P_qe_* for FtsQ was equal to about 93%. The related quantity P_SS_ is equal to the ratio between the numbers of RNC partitioning into the bilayer and RNCs that did not. P_SS_ allows calculating ΔG_S_:

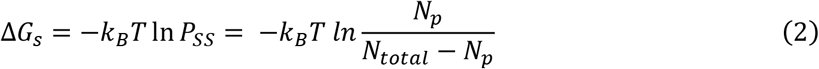

The resulting Δ*G_s_* = −1.5 kcal/mol suggests that the segment is less hydrophobic as predicted by the position-dependent bio-scale (13) and by the physicochemical scale (9) (Table 2).

The underestimation is due to the leakage current characteristic for stalled translocation intermediates. For example, we have previously demonstrated that the translocation intermediate proOmpA produces a leakage current *I_L_* of about 1 pA through the closed SecYEG channel. *I_L_* increases to about 3 pA for the intermediate with plugless SecY (30). Our current records reveal *I_L_* ≈ 2 pA for SecYEG-RNC(FtsQ). Eq. 1 can be corrected for the leakage as:

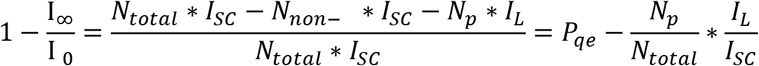

Here, *I_SC_* is the current through the single open translocation intermediate, *N_non–p_* is the number of translocation intermediates with non-partitioned nascent chains. This correction will be most prominent for hydrophobic sequences, when 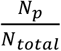 is close to unity. Then, 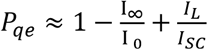. Knowing that 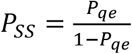 and that 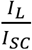 is about 0.04, we arrive at the corrected ΔG_s_ values given in Table 2 in parentheses. For moderately hydrophobic sequences, this correction is insignificant.

Our electrophysiological approach allows monitoring how *P*_qe_ approaches its steady-state value. To quantify the process, we define the time-dependent partitioning coefficient *P_dyn_*(*t*):

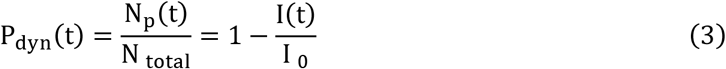

where *P*_dyn_(∞) = *P*_qe_. *I*(*t*) is the current at a given time *t*. *P_dyn_* significantly deviates from *P_qe_* for short observation times.

### The steady-state probability of SecYEG-RNC complex closure is voltage-dependent

We reconstituted the translocon-RNC complexes KvAP 3R, a mutant KvAP-voltage sensor helix in which one of the four arginines was replaced by an alanine. Large membrane potentials, ψ = −110 mV, led to rapid channel closure, indicating membrane integration of the nascent chain (Fig. 4). In contrast, small values of transmembrane potential, ψ = −15 mV, did not close the channel. Application of intermediate ψ values (ψ = −65 mV) slowly reduced ion channel activity.

**Figure 4.**
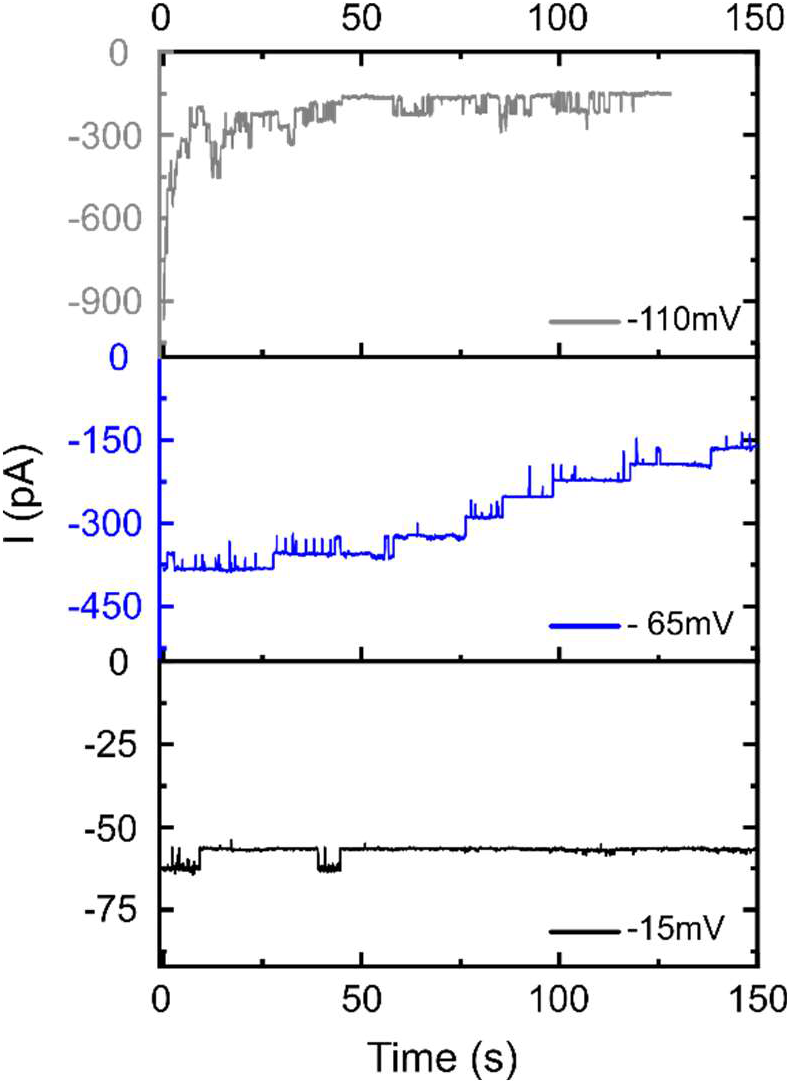
The voltage-gated closing of SecYEG is a prerequisite for nascent chain partitioning into the membrane. Online partitioning of KvAP-based TMD with three arginines (KvAP 3R, see Table 1) from translocation intermediates at physiologically high (−110 mV), middle (−65 mV), and low (−15 mV) transmembrane potential ψ. The apparent equilibrium partitioning for each trace was calculated using Eq. 1. It was 0.8 for ψ = −110 mV, 0.6 for −65 mV, and 0.1 for −15 mV

The observed voltage effect agrees with earlier observations made with RNCs based on a β-barrel preprotein from the outer bacterial membrane (28). Up to a certain voltage threshold, the steady-state closure probability and the time required to reach the steady state are voltage-dependent. Thus to determine *P*_qe_ or *P*_SS_, application of high voltage is required. This condition represents an experimental challenge as solvent-depleted planar bilayers with reconstituted SecYEG– RNC complexes may become unstable for ψ < −100 mV.

Small transmembrane potentials allowed extracting the total amount of locked translocation intermediates, *N*_total_. Absolute ψ values higher than 100 mV revealed the steady-state (quasi-equilibrium) probability of membrane helix partitioning, *P_qe_*. Without the transmembrane helix exiting into the bilayer, the lateral gate would have been unable to close.

### Determing ΔG_s_ in quasi-equilibrium

The third RNC construct contained the KvAP’s wild-type sensor helix TM4. To distinguish the four arginines bearing wild-type segment from its variants, we labeled the nascent chain as KvAP 4R (see Table 1). To enable quantitative analysis, we added up all available individual traces (such as the traces in Fig. 5A) to yield a cumulative trace (Fig. 5B). Extracting *P_dyn_* from the transmembrane current (Eq. 3) allowed calculation of the characteristic partitioning time *τ*, according to Eq. 4:

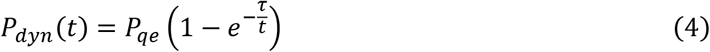

*τ* for the 19 amino acid-long segment was equal to 12.4 s. At high ψ values, the resulting *P_eq_* was equal to 37 % (Fig. 5A, B). Accordingly, *ΔG_s_* amounted to 0.3 kcal/mol. This perfectly agrees with the value predicted by the position-dependent biological scale (Table 2).

**Figure 5.**
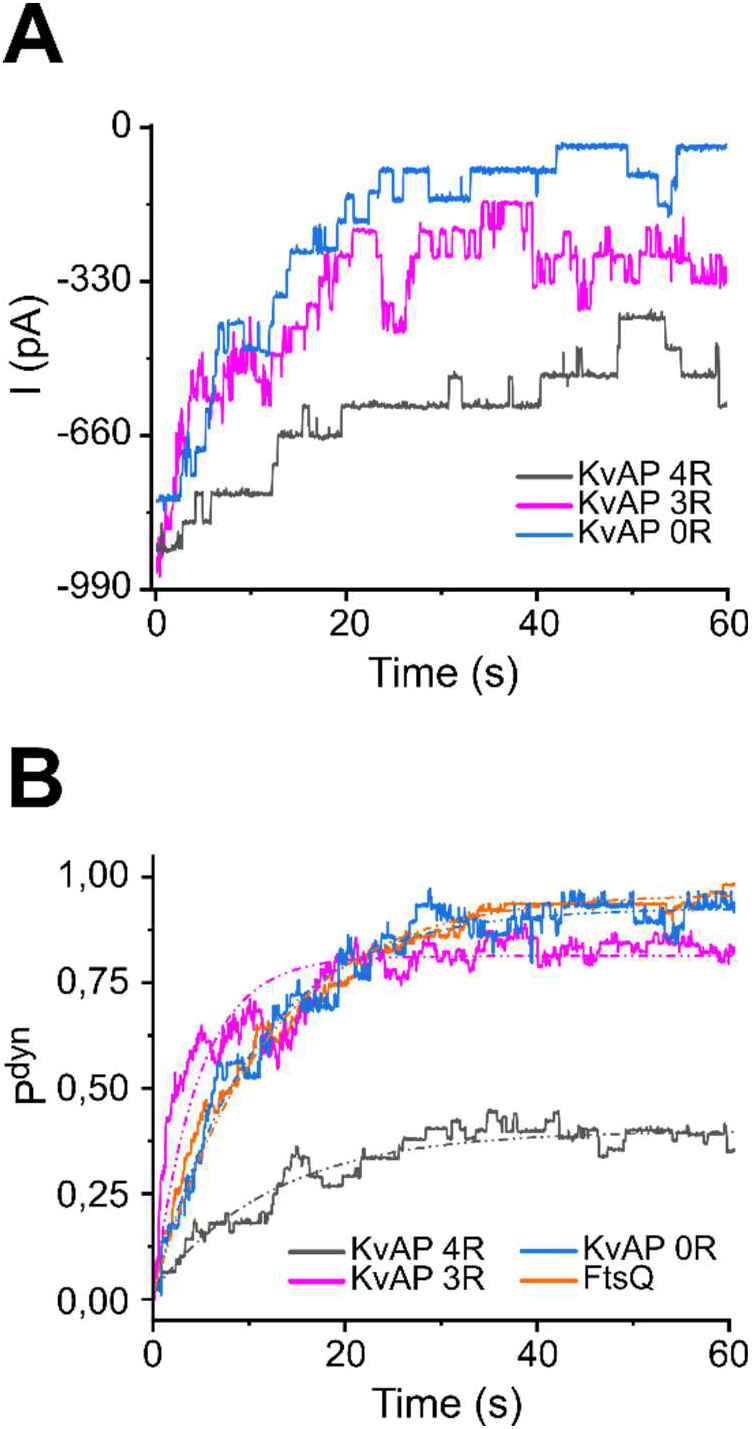
Online partitioning. **A:** Online partitioning of RNC containing a KvAP-based TMD from the locked translocation intermediate SecYEG-RNC, where KvAP 4R corresponds to the original S4 segment of KvAP with four arginines, KvAP 3R – to the S4 variant with one of the middle arginines substituted with alanine, KvAP 0R – to the S4 variant where all four arginines were substituted with alanines (see Table 1). As in A, the traces represent the experimentally observed voltage-triggered partitioning events. **B:** Cumulative partitioning kinetics for all KvAP-based RNCs. We calculated P^dyn^ from cumulative current traces (representative traces shown in A). Fitting a single-exponential function (Eq. 4, solid lines, see also Table 2) to Pdyn yielded the characteristic integration time. The color code is the same as in A. Partitioning of an FtsQ-based nascent chain is shown for comparison (green, data taken from Fig. 3). Cumulative traces were collected from the total number of translocation intermediates N = 55 for FtsQ-based TMDs, N = 47 for 4R, N = 45 for 3R and N = 28 for 0R for KvAP-based TMDs.

### The charge effect on the partitioning probability

From membrane integration experiments with microsomes (24), it is known that the removal of the central arginine has a profound effect on the partitioning of the TM4 segment of KvAP (see Table 2). We compared constructs analogous to KvAP 4R-RNC, where one of the central arginines (KvAP 3R-RNC) or all arginines (KvAP 0R-RNC) were substituted with alanines (Fig. 5). The respective Δ*G_s_* values amounted to −0.8 kcal/mol for KvAP 3R-RNC and −1.3 kcal/mol for KvAP 0R-RNC. The Δ*G_s_* for the KvAP 3R sequence is in good agreement with the value predicted by the position-dependent bio-scale. However, as in the case of FtsQ-RNC, our Δ*G_s_* underestimates the energy gain in the case of strongly hydrophobic sequences (see Table 2).

## Discussion

We determined the steady-state probability of polypeptide membrane integration in a minimalistic system consisting of a reconstituted bacterial SecYEG-RNC complex. Since the 19 amino acid-long transmembrane segments had ample time for sampling between the different environments of the aqueous translocon interior and the hydrophobic membrane core, this probability reflects the energetic expense for their membrane integration in quasi-equilibrium. Remarkably, our ΔG_s_ is in reasonable agreement with the ΔG_s_b_ derived from H-segment insertion experiments previously carried out with mammalian microsomes, i.e., with the location-corrected biological hydrophobicity scale (13). For example, our ΔG_s_ of 0.3 kcal/mol for membrane integration of KvAP’s fourth transmembrane helix matched the ΔG_s_b_ reported for its insertion in the endoplasmic reticulum. The equality of both free energy values indicates that translocon’s decision for membrane integration versus secretion strictly follows thermodynamic principles and that kinetic factors play a negligible role. The result agrees with the reported direct proportionality of amino acid’s contribution to the apparent free energy of membrane insertion and the non-polar accessible surface area of its side chain (40).

Our conclusion is also in line with the observed insertion kinetics. For FtsQ’s transmembrane segment, we found an insertion rate of 0.2 amino acids per second. This rate is comparable with previously observed secretion rates and measured amino acid translation rates at the ribosome. Longer integration times recorded upon restricting polypeptide movement by allowing calmodulin to bind to a tag on the nascent chain in the receiving compartment only reflect an artificially increased activation barrier for membrane integration. It is very likely that the large proteinaceous anchor decreased polypeptide mobility. In consequence, every move of the segment between the translocon and the membrane may take more time. Thus, the nascent chain bound to calmodulin could take more time to perform the minimum 100 sampling motions that the molecular dynamics simulations suggest it would need to approach the correct equilibrium probability of membrane integration (27).

Our calmodulin binding experiment displays interesting parallels with chaperone binding experiments carried out in yeast (25). In contrast to our reconstitution experiments, the yeast assay did not allow determining integration rates, but it revealed changes in integration probability depending on (i) chaperone binding in the receiving compartment or (ii) misfolding of upstream segments. Both modifications altered the environmental hydrophobicity probed by the nascent chain. In case (i), the chaperone removed a potential interaction partner for the emerging transmembrane segment, and in case (ii), the misfolded structure presented a hydrophobic surface capable of attracting the hydrophobic segment otherwise destined for membrane integration. Thus, this study does not contradict our conclusion that thermodynamics instead of kinetic factors govern membrane integration.

Our electrophysiological assay may underestimate ΔG_s_ for hydrophobic sequences (FtsQ-RNC and KvAP 0R-RNC), i.e., the position-dependent biological scale predicts higher ΔG_s_b_ values (see Table 2). The deviation would be caused by the leakage current passing through an uncertain number of closed intermediates. However, the smaller hydrophobicity range may also be due to a compression of the hydrophobicity scale caused by the prokaryotic translocon relative to the mammalian translocon. Such alterations in translocation thermodynamics are known from yeast (14). Moreover, they would reflect corresponding changes in protein folding thermodynamics (41).

Our results show quasi-equilibrium partitioning of TM segments in a minimal system where SecYEG lacks accessory proteins. They are reasonably close to the prediction by the position-dependent biological hydrophobicity scale obtained in dog pancreas rough microsomes (13). We thus demonstrate that TM insertion *in vivo* may well satisfy thermodynamic equilibrium.

## References

1. M. Spiess, T. Junne, M. Janoschke, Membrane Protein Integration and Topogenesis at the ER. Protein J. 10.1007/s10930-019-09827-6 (2019).

2. T. A. Rapoport, L. Li, E. Park, Structural and Mechanistic Insights into Protein Translocation. Annual review of cell and developmental biology 33, 369–390 (2017).

3. F. Cymer, G. von Heijne, S. H. White, Mechanisms of integral membrane protein insertion and folding. J. Mol. Biol. 427, 999–1022 (2015).

4. A. Elazar et al., Mutational scanning reveals the determinants of protein insertion and association energetics in the plasma membrane. eLife 5, e12125 (2016).

5. D. G. Knyazev, R. Kuttner, M. Zimmermann, E. Sobakinskaya, P. Pohl, Driving Forces of Translocation Through Bacterial Translocon SecYEG. The Journal of membrane biology 10.1007/s00232-017-0012-9 (2018).

6. J. Koehler, N. Woetzel, R. Staritzbichler, C. R. Sanders, J. Meiler, A unified hydrophobicity scale for multispan membrane proteins. Proteins: Structure, Function, and Bioinformatics 76, 13–29 (2009).

7. J. L. MacCallum, D. P. Tieleman, Hydrophobicity scales: a thermodynamic looking glass into lipid–protein interactions. Trends in Biochemical Sciences 36, 653–662 (2011).

8. A. L. Lomize, I. D. Pogozheva, M. A. Lomize, H. I. Mosberg, Positioning of proteins in membranes: a computational approach. Protein Sci. 15, 1318–1333 (2006).

9. W. C. Wimley, S. H. White, Experimentally determined hydrophobicity scale for proteins at membrane interfaces. Nature Structural & Molecular Biology 3, 842–848 (1996).

10. T. Hessa et al., Recognition of transmembrane helices by the endoplasmic reticulum translocon. Nature 433, 377–381 (2005).

11. S. Capponi, M. Heyden, A. N. Bondar, D. J. Tobias, S. H. White, Anomalous behavior of water inside the SecY translocon. Proc. Natl. Acad. Sci. U.S.A 112, 9016–9021 (2015).

12. M.-C. Bellissent-Funel et al., Water Determines the Structure and Dynamics of Proteins. Chem. Rev. 116, 7673–7697 (2016).

13. T. Hessa et al., Molecular code for transmembrane-helix recognition by the Sec61 translocon. Nature 450, 1026–1030 (2007).

14. T. Hessa, J. H. Reithinger, G. von Heijne, H. Kim, Analysis of transmembrane helix integration in the endoplasmic reticulum in S. cerevisiae. J. Mol. Biol. 386, 1222–1228 (2009).

15. T. Junne, L. Kocik, M. Spiess, The hydrophobic core of the Sec61 translocon defines the hydrophobicity threshold for membrane integration. Mol Biol Cell 21, 1662–1670 (2010).

16. A. Missner, P. Pohl, 110 years of the Meyer-Overton rule: predicting membrane permeability of gases and other small compounds. Chemphyschem 10, 1405–1414 (2009).

17. C. Hannesschlaeger, A. Horner, P. Pohl, Intrinsic Membrane Permeability to Small Molecules. Chemical reviews 119, 5922–5953 (2019).

18. F. Zocher, D. van der Spoel, P. Pohl, J. S. Hub, Local partition coefficients govern solute permeability of cholesterol-containing membranes. Biophys. J. 105, 2760–2770 (2013).

19. A. Ebert, C. Hannesschlaeger, K.-U. Goss, P. Pohl, Passive Permeability of Planar Lipid Bilayers to Organic Anions. Biophys. J. 115, 1931–1941 (2018).

20. B. Chanda, O. K. Asamoah, R. Blunck, B. Roux, F. Bezanilla, Gating charge displacement in voltage-gated ion channels involves limited transmembrane movement. Nature 436, 852–856 (2005).

21. D. J. Posson, P. Ge, C. Miller, F. Bezanilla, P. R. Selvin, Small vertical movement of a K+ channel voltage sensor measured with luminescence energy transfer. Nature 436, 848–851 (2005).

22. Y. Jiang, V. Ruta, J. Chen, A. Lee, R. MacKinnon, The principle of gating charge movement in a voltage-dependent K+ channel. Nature 423, 42–48 (2003).

23. C. A. Ahern, R. Horn, Stirring up controversy with a voltage sensor paddle. Trends Neurosci. 27, 303–307 (2004).

24. T. Hessa, S. H. White, G. Heijne, Membrane insertion of a potassium-channel voltage sensor. Science (New York, N.Y.) 307, 1427 (2005).

25. M. Janoschke et al., Efficient integration of transmembrane domains depends on the folding properties of the upstream sequences. Proc Natl Acad Sci U S A 118 (2021).

26. E. V. Schow et al., Arginine in membranes: the connection between molecular dynamics simulations and translocon-mediated insertion experiments. J Membr. Biol 239, 35–48 (2011).

27. J. C. Gumbart, I. Teo, B. Roux, K. Schulten, Reconciling the Roles of Kinetic and Thermodynamic Factors in Membrane–Protein Insertion. Journal of the American Chemical Society 135, 2291–2297 (2013).

28. D. G. Knyazev et al., Voltage Sensing in Bacterial Protein Translocation. Biomolecules 10(2020).

29. D. G. Knyazev et al., The bacterial translocon SecYEG opens upon ribosome binding. The Journal of biological chemistry 288, 17941–17946 (2013).

30. D. G. Knyazev, L. Winter, B. W. Bauer, C. Siligan, P. Pohl, Ion conductivity of the bacterial translocation channel SecYEG engaged in translocation. The Journal of biological chemistry 289, 24611–24616 (2014).

31. I. Sachelaru et al., YidC and SecYEG form a heterotetrameric protein translocation channel. Scientific reports 7, 101 (2017).

32. J. Ederth, C. S. Mandava, S. Dasgupta, S. Sanyal, A single-step method for purification of active His-tagged ribosomes from a genetically engineered Escherichia coli. Nucleic Acids Res 37 (2009).

33. S. M. Saparov et al., Determining the Conductance of the SecY Protein Translocation Channel for Small Molecules. Molecular Cell 26, 501–509 (2007).

34. T. Hoomann, N. Jahnke, A. Horner, S. Keller, P. Pohl, Filter gate closure inhibits ion but not water transport through potassium channels. Proceedings of the National Academy of Sciences of the United States of America 110, 10842–10847 (2013).

35. D. J. Woodbury, J. E. Hall, Role of channels in the fusion of vesicles with a planar bilayer. Biophys. J. 54, 1053–1063 (1988).

36. D. J. Woodbury, C. Miller, Nystatin-induced liposome fusion. a versatile approach to ion channel reconstitution into planar bilayers. Biophys. J. 58, 833–839 (1990).

37. F. C. Liang, U. K. Bageshwar, S. M. Musser, Bacterial Sec protein transport is rate-limited by precursor length: a single turnover study. Mol Biol Cell 20, 4256–4266 (2009).

38. J. F. Ross, M. Orlowski, Growth-rate-dependent adjustment of ribosome function in chemostat-grown cells of the fungus Mucor racemosus. J Bacteriol 149, 650–653 (1982).

39. D. G. Dalbow, R. Young, Synthesis time of beta-galactosidase in Escherichia coli B/r as a function of growth rate. Biochem J 150, 13–20 (1975).

40. K. Ojemalm et al., Apolar surface area determines the efficiency of translocon-mediated membrane-protein integration into the endoplasmic reticulum. Proc. Natl. Acad. Sci. U.S.A 108, E359–364 (2011).

41. U. Bastolla, A. Moya, E. Viguera, R. C. H. J. van Ham, Genomic Determinants of Protein Folding Thermodynamics in Prokaryotic Organisms. J. Mol. Biol. 343, 1451–1466 (2004).

