## Supplementary Information for "Steady-state polypeptide transfer from the translocon to the membrane"

### Purification of RNC complexes.

***FTSQ101 CBP TEV SecM RNCs:***

The sequence of the nascent chain with the corresponding color code below:

MSQAALNTRN SEEEVSSRRN NGTRLAGGGK RRWKKNFIAV SAANRFKKIS SSGALGGLF LLTVLTTVLV SGWVVLGWME DAQRLPLSKL VLTGERHYTR NDDIRQSILA LGEPGTFMT QDVNIIQTQI EQQNENLYFQ SPSEKGYRID YAHFTPQAKF STPVWISQAQ GIRAGP*

FtsQ

Spacing

Calmodulin Binding Protein (CBP)

TEV Cleavage Site

SecM stalling Sequence

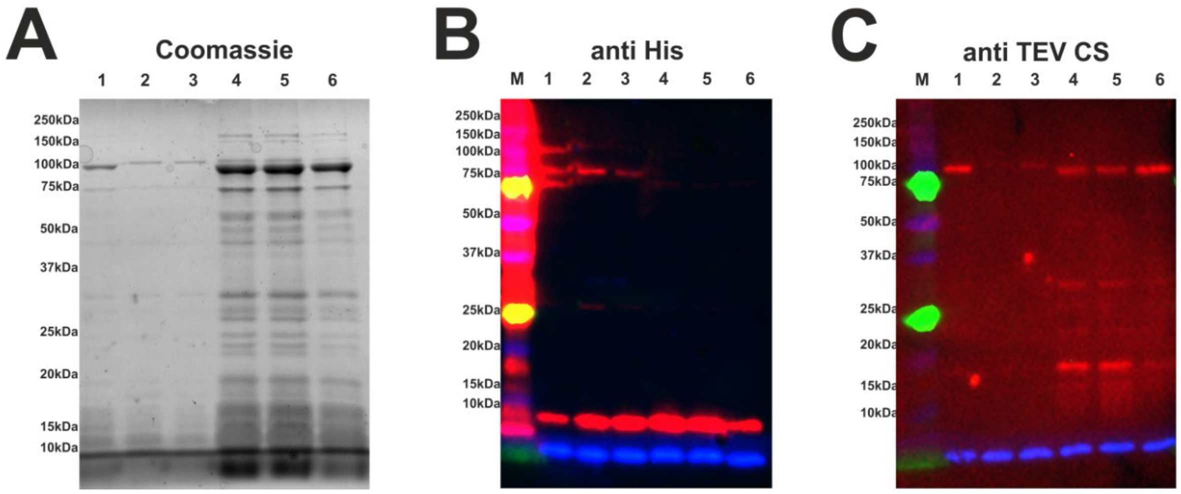


**Figure S1: Purification of FTSQ101 CBP TEV SecM RNCs.** Six sequential elutions with the highest content of the protein of interest (lanes 1-6) after sequential affinity chromatography steps (B, C) of the tandem purification (see Methods) are shown. The general protein composition of the samples was analyzed with Coomassie staining (A). Ribosomal protein L12 as well as the nascent chain were specifically detected with anti-His (B) and anti-TEV-site (C) antibodies, respectively. Western blots were detected with a BioRad geldoc system and are colored as follows: immune histochemistry signal of the HRP-conjugated primary anti-His AB (Miltenyi Biotech) or the HRP conjugated secondary anti-rabbit AB (SigmaAldrich) in case of non-conjugated primary anti-TEV AB (ThermoFisher) (both were detected in Cy 2 channel, shown in red); far-red fluorescence of the Coomassie running front (Cy5 channel, shown in blue); red fluorescence of specific marker bands (Cy3 channel, shown in green). Marker: Precision Plus Dual Color (BioRad, #1610374).

The nascent chain cannot be discerned (expected size: 19 kDa) in the Coomassie band pattern (Figure S1-A). Nonetheless, detection via western blot using anti-TEV-site antibodies reveals the respective bands (Figure 1-C lanes 4-6). Furthermore, it can be observed that the respective signal with the correct size only emerges within later elutions. In Figure S2B, a faint band at about 100 kDa was detected in earlier elutions. This is due to the fact that chelators of bivalent ions such as EGTA may help in the dissociation of the ribosome. As the first elutions of the sample show a significant amount of L12 protein (Figure S1A, lanes 1-3), it can be assumed that dissolved ribosomes were eluted. Consequently, only elution 4-6 were pooled and used for the first pilot experiments. To mitigate the presence of dissolved ribosomes, we centrifuged RNC samples at 100,000 g in order to pellet the much larger full-size ribosomes together with bound RNCs.

**FtsQ101 CBP KVAP S4 4R Tev SecM**

The sequence of the nascent chain with the corresponding color code below:

MSQAALNTRN SEEEVSSRRN NGTRLAGGGK RRWKKNFIAV SAANRFKKIS SSGALGGFRL VRLLRFLRIL LIISDAQRLP LSKLVLTGER HYTRNDDIRQ SILALGEPGT FMTQDVNIIQ TQIEQQNENL YFQSPSEKGY RIDYAHFTPQ AKFSTPVWIS QAQGIRAGP*

KvaP H4

FtsQ

Spacing

Calmodulin Binding Protein (CBP)

TEV Cleavage Site

SecM stalling Sequence

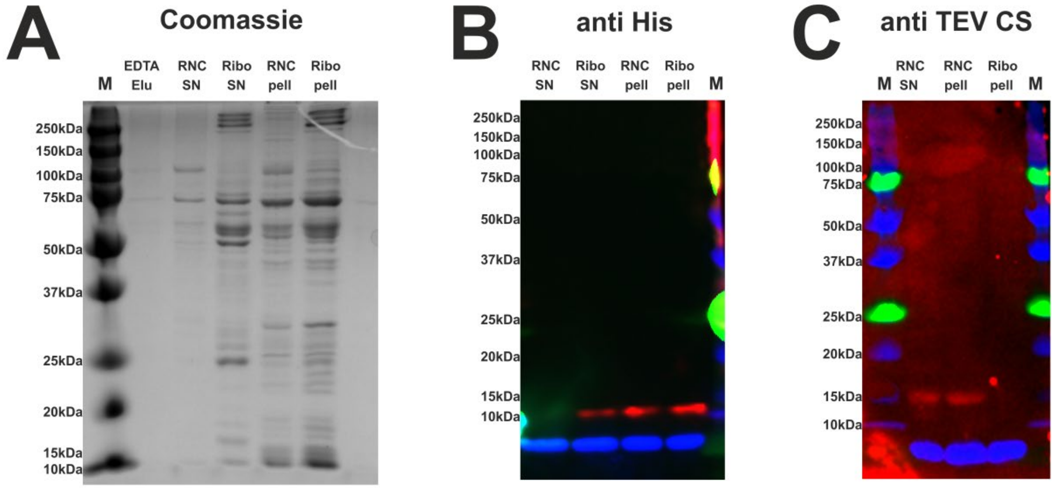


**Figure S2: Purification of FtsQ101 CBP KVAP S4 4R Tev SecM RNCs.** Coomassie staining (A), as well as western blots with antibodies specific for either His-tag (B) or TEV cleavage site (C), were used to analyze RNC samples. The presented here samples are pooled together elutions (see Fig. S1). The lane termed “Ribo pell” represents the sample derived from the resuspended pellet (after ultracentrifugation) of the unbound calmodulin agarose flowthrough, with “Ribo SN” being the supernatant of the same centrifugation step. Similarly, “RNC pell” and “RNC SN” represent the resuspended pellet and supernatant of the pooled and ultra-centrifuged elutions of the calmodulin agarose purification step. “EDTA Elu” is the final elution with 50 mM EDTA to elute all remaining proteins and was not subjected to centrifugation. Western blots were detected with a BioRad geldoc system and are colored as explained in Fig. S1.

The purification of KVAP 4R RNCs (lanes termed “RNC pell”) delivered similar results compared to the FtsQ-based RNCs (Figure S1) when examining the Coomassie (Figure S2-A), anti-His (Figure S2B) and anti-TEV-site western blots (Figure S2C). A ribosome-like band pattern is visible in the Coomassie-stained gel as well as specific detection for both L12 and the nascent chain.

To further test the efficacy of the purification, the unbound proteins from the calmodulin agarose flowthrough, which should mainly contain ribosomes, were loaded as well. When this sample was precipitated at 100,000 g for 3 h and subsequently resuspended, it exhibited a ribosomal band pattern after Coomassie staining (Figure S2-A; “Ribo pell”). Furthermore, ribosomal L12 protein was detectable (Figure S2B; “Ribo pell”), while no nascent chains could be observed (Figure S2C; “Ribo pell”). Supernatant samples after centrifugation show similar results, with lower protein amounts (Figure 2; “Ribo SN”). On the other hand, the supernatant sample originating from the calmodulin elution step showed a degree of unbound nascent chain (Figure S2C, “RNC SN”), while no significant amount of ribosomes could be detected (Figure S2A and -B; “RNC SN”). In addition, after successful elution with EGTA, another elution step with 50 mM EDTA was performed, which did not elute any significant amount of protein (Figure S2A, “EDTA Elu”), proving the efficacy of the elution protocol.

**<u>Determination of SecYEG reconstitution efficiency into lipid vesicles.</u>**

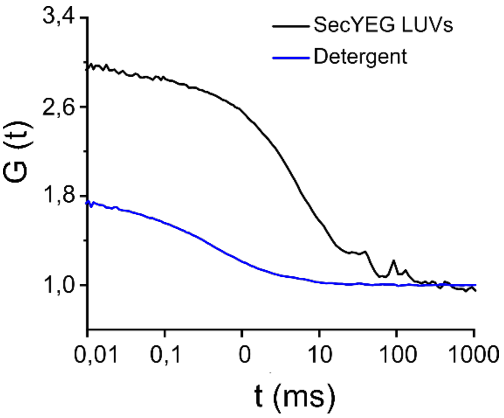


**Figure S3: Number of SecYEG complexes per vesicle measured by fluorescence correlation spectroscopy.** Autocorrelation curves of SecYEG-vesicles in 50 mM K-HEPES, pH 7.5 (black line), and in a detergent mix 2% octyl glucoside and 3% deoxy big CHAP. The number of particles per confocal volume in each case is obtained by fitting the autocorrelation curves with the equation $G(t) = 1 + \frac{1}{n(1+\frac{t}{t_D})}$, where n is the number of fluorescent particles in the confocal volume, $\tau_D$ is a residence time of the fluorescent particle in confocal volume. The number of particles increased three-fold after the solubilization of proteoliposomes in the detergent mix.
